# The laminar pattern of proprioceptive activation in human primary motor cortex

**DOI:** 10.1101/2023.10.29.564658

**Authors:** Lasse Knudsen, Fanhua Guo, Jiepin Huang, Jakob U. Blicher, Torben E. Lund, Yan Zhou, Peng Zhang, Yan Yang

## Abstract

The primary motor cortex (M1) has traditionally been viewed as a motor output generator. However, its vital role in proprioceptive somatosensation is increasingly being recognized. Yet, our understanding of proprioceptive somatosensation in M1 at the laminar scale is limited, largely due to methodological challenges. Empirical findings in primates and rodents suggest a pronounced role of superficial cortical layers, but the involvement of deep layers has yet to be examined in humans. Submillimeter resolution fMRI has emerged in recent years, paving the way for the study of layer-dependent activity in humans (laminar fMRI). In the present study, laminar fMRI was employed to investigate the laminar pattern of proprioceptive somatosensation on M1 deep layer activation using passive finger movements as proprioceptive stimulation. Significant M1 deep layer activation was observed in response to proprioceptive stimulation across 10 healthy subjects using vascular space occupancy (VASO) at 7T. For further validation, two additional subjects were scanned using a balanced steady-state free precession (bSSFP) sequence with ultrahigh (0.3 mm) in-plane resolution, yielding converging results. These results were interpreted in the light of previous laminar fMRI studies and the active inference account of motor control. We suggest that a considerable proportion of M1 deep layer activation is due to proprioceptive influence and that deep layers of M1 constitute a key component in proprioceptive circuits.

## 1. Introduction

The primary motor cortex’s main function is to generate corticospinal output to facilitate voluntary movement. It relies heavily on contextual information from other brain regions associated with, for instance, limb position, limb kinematics, motor planning, and motor intent. Our present understanding of the circuitry underlying information passing in M1 is primarily derived from tracer studies in rodents and nonhuman primates (Figure 1A). Briefly, somatosensory input has consistently been reported to terminate in superficial depths of M1 (layers II-III and layer Va) (Adams et al., 2013; Hooks, 2017; Mao et al., 2011; Petrof et al., 2015; Shipp, 2005), and the reciprocal connection to the primary somatosensory cortex (S1) also originates from these layers (Hooks, 2017; Mao et al., 2011). Motor-related inputs from frontal and secondary motor areas may terminate more widely across the full cortical depth, but appear to favor superficial layers (Adams et al., 2013; Barbas & Pandya, 1987; Hooks, 2017; Ninomiya et al., 2019; Watanabe-Sawaguchi et al., 1991). Importantly, these inputs are transmitted from superficial depths to pyramidal cells in layer V via strong interlaminar connections where they influence corticospinal and corticothalamic outputs (Bastos et al., 2012; Shipp, 2016; Weiler et al., 2008). The majority of corticofugal output cells are located in deep layers Vb (corticospinal), and VI (corticothalamic). Existence of a comparable input/output structure in human M1 is currently unknown, primarily due to a lack of human-relevant methodology operating at the scale of cortical layers.

**Figure 1.**
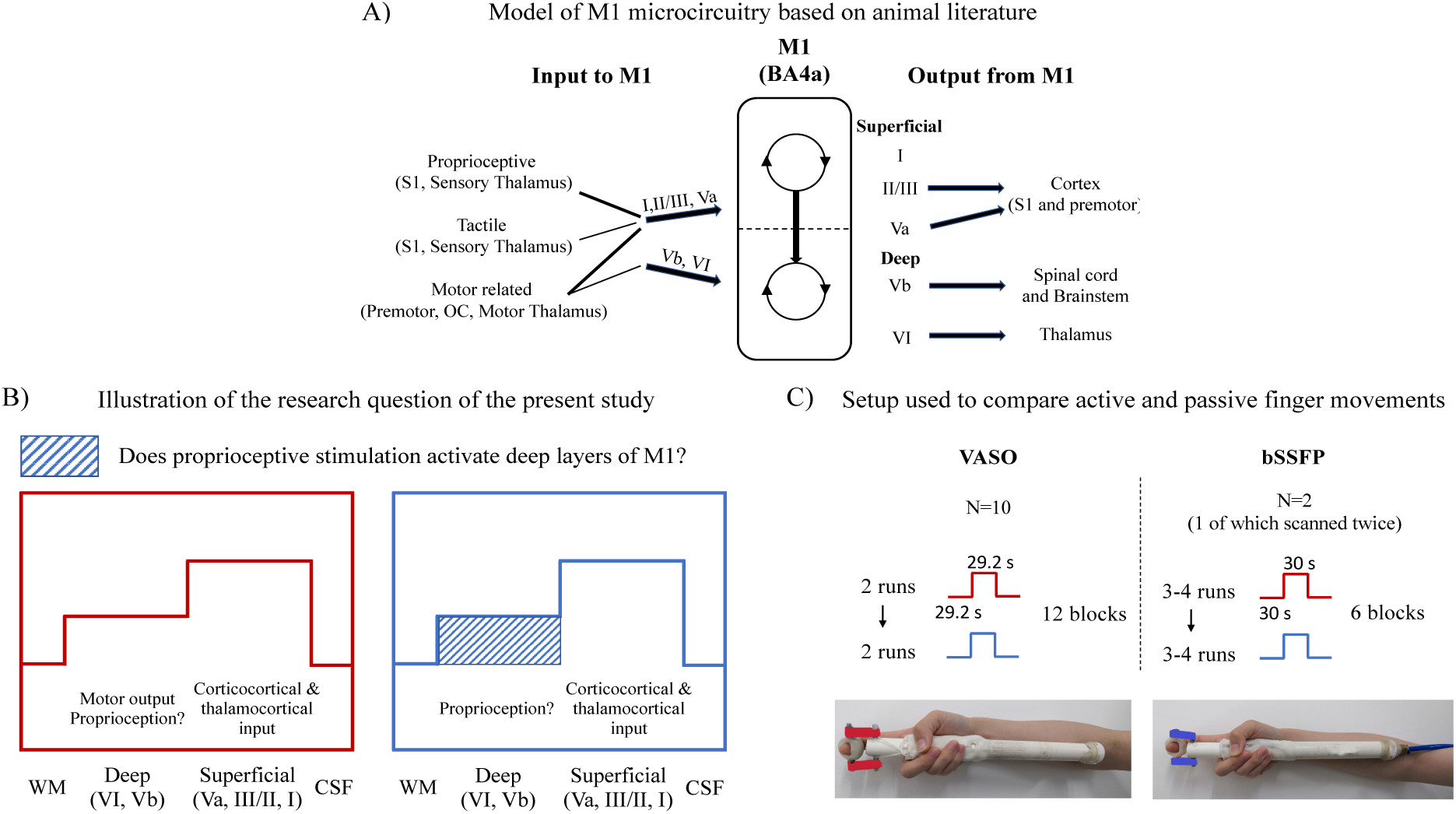
Study overview. **A)** Model of possible M1 microcircuitry, based on animal literature (references listed in main text). The majority of input to M1 arrives in superficial layers, and the majority of output (corticofugal) originates from deep layers. The strong intra-columnar projection from superficial to deep layers is consistent with the view that preprocessed information from superficial layers is integrated by deep layers before the resulting output is directed to, e.g., the spinal cord (Bastos et al., 2012). Superficial and deep layers can thus be roughly segregated into input and output layers, respectively. **B)** Illustration of the research questions of the present study. Previous laminar fMRI studies found robust M1 deep layer activation during active finger movements, interpreted to primarily reflect motor output (see main text). Here, we test whether passive movement (proprioceptive stimulation, no motor output) also activates deep layers of M1. **C)** Illustration of the methodological setup used to assess these questions. Two different fMRI sequences were used with a complimentary set of strengths and weaknesses, i.e., VASO and bSSFP. Both experiments included block-designed active and passive finger movement paradigms performed in different runs (order was counterbalanced across subjects). The finger was moved voluntarily by the subject’s own muscle force during active finger movements (red device). For passive movements (blue device), subjects would relax while their finger was moved by a fMRI compatible pneumatic muscle driven by air pressure (Lolli et al., 2019).

The advent of laminar fMRI (Bandettini et al., 2021; Lawrence et al., 2019; Norris & Polimeni, 2019) has enabled assessment of the layer-dependent functional organization in humans, underpinning prominent theories of brain function, such as predictive coding (Rao & Ballard, 1999; Stephan et al., 2019). Laminar fMRI was employed by Huber and colleagues (2017) to compare laminar activation of the M1 hand representation in response to tasks with varying degrees of motor and somatosensory involvement. Active finger movements elicited a double peak response pattern with signal increases in both superficial and deep layers, which was replicated in several subsequent studies (Beckett et al., 2020; Chai et al., 2020; Guidi et al., 2020; Persichetti et al., 2020; Shao et al., 2021). Moreover, tactile stimulation (Huber et al., 2017; Shao et al., 2021) and imagined finger movements (Persichetti et al., 2020), both of which were thought to lack corticospinal motor output, were found to predominantly activate superficial layers. Informed by the animal literature (see Figure 1A), the general interpretation across these studies (also see Dumoulin, 2017; Larkum et al., 2018; McColgan et al., 2020) has been that superficial peak activation mainly reflects somatosensory and premotor input while the deep peak reflects motor output.

Nevertheless, due to a paucity of studies employing a direct proprioceptive control condition, the pattern of laminar activation associated with proprioceptive somatosensation is currently unknown. Although tactile somatosensory processing appears to predominantly engage the superficial layers of M1, proprioceptive somatosensory processing may have a different structure. First, M1 is no longer considered solely a motor area, and its central function in somatosensory processing is well-established (Hatsopoulos & Suminski, 2011; Naito, 2004, 2011). According to the influential active inference account of motor control (Friston, 2010), M1 is part of a somatomotor hierarchy generating motor action and kinesthetic perception. It can be viewed as a sensorimotor inference machine aiming to minimize errors associated with proprioceptive predictions (Adams et al., 2013; Friston, 2011; Shipp et al., 2013). Within this framework, the output of M1 to the spinal cord and to hierarchically lower somatosensory areas are regarded as proprioceptive predictions rather than motor commands and efference copies, respectively. According to the canonical microcircuit (Bastos et al., 2012), proprioceptive prediction errors from somatosensory areas terminate in the superficial layers of M1, whereas descending proprioceptive predictions from M1 to somatosensory areas and the spinal cord originate in layers V and VI. Consequently, proprioception is expected to activate both superficial and deep layers. Second, the lack of robust deep layer activation during tactile somatosensory stimulation might be partially explained by the fact that signals were extracted from a region with relatively few tactile responsive cells (Rathelot & Strick, 2009), namely the phylogenetically older and upper part of M1 (BA4a) (Huber et al., 2017; Shao et al., 2021). In contrast, proprioceptive responsive cells are particularly concentrated in this region (Stepniewska et al., 1993; Strick & Preston, 1982; Tanji & Wise, 1981). Last, in human PET and fMRI studies, proprioceptive stimulation induced by passive movements consistently activated M1 (Blatow et al., 2011; Ciccarelli et al., 2005; Guzzetta et al., 2007; Jaeger et al., 2014; Mehta et al., 2012; Weiller et al., 1996). Part of this signal could plausibly represent deep layer activation as indicated by electrophysiological non-human primate studies in which pyramidal tract cells activated in response to proprioceptive stimulation (Lemon & Porter, 1976; Wiesendanger, 1973).

In the present study, we aim to determine the laminar response of human M1 to proprioceptive stimulation. Specifically, we used 7T laminar fMRI with VASO (Huber et al., 2014) and bSSFP (Miller, 2012; Scheffler et al., 2018) sequences to examine whether passive finger movements (proprioceptive stimulation) elicit deep layer responses in the hand knob of contralateral M1 (Figure 1B). Addressing this basic neuroscience question may advance our understanding of laminar function in human M1, and progress the development and validation of theories on motor control and somatosensory perception.

## 2. Methods

### 2.1 Subjects

A total of 12 healthy volunteers (5 females) with a mean age of 25 years (range 23-29 years) participated in the study. Before participation, all subjects provided written consent and were carefully informed about the procedures, approved by the institutional review board of the Institute of Biophysics, Chinese Academy of Sciences.

### 2.2 Active and passive movement devices

Passive finger tapping was performed using a fMRI-compatible device inspired by (Lolli et al., 2019) (Figure 1C). It was driven by a fluidic muscle (DMSP type, diameter 10 mm, length of the contracting part 150 mm; Festo AG & Co, Esslingen, Germany), which shortens upon increased air pressure and extends towards its initial position upon pressure release. Air supply was controlled via a pre-programmed electrical valve to obtain the desired frequency and amplitude. An identical device was used for active tapping except it did not contain any muscle (Figure 1C). The design goal was to make active and passive movements as similar as possible to standardize sensory input. A custom-made hand print was positioned around each device for each subject to standardize the amplitude of vertical movement across conditions while enabling subjects to switch devices within the scanner. A soft padding was inserted inside the active device in order to match impact forces between the movable part (to which the finger is attached) and the outer tube across conditions.

### 2.3 Imaging protocol

Imaging was performed using a MAGNETOM 7T MRI scanner (Siemens Healthineers, Erlangen, Germany) equipped with a standard 32Ch-receive head coil (NOVA Medical, Wilmington, MA, USA) and a SC72 body gradient coil for spatial encoding. Anatomical T1-weighted images were acquired with a MP2RAGE sequence (Marques et al., 2010) and parameters: TR = 4000 ms, TE = 3.05 ms, voxel size = 0.7 mm isotropic, FOV = 224 mm × 224 mm × 179.2 mm, GRAPPA acceleration factor = 3, TI1 = 750 ms with flip angle = 4 deg and TI2 = 2500 ms with flip angle = 5 deg, phase and slice partial Fourier factor = 7/8, Bandwidth = 240 Hz/px). Two different sequences were used in different sessions for functional acquisitions, i.e., cerebral blood volume (CBV)-weighted 3D-EPI SS-SI-VASO (Huber et al., 2014), and 2D passband single-slice bSSFP (Miller, 2012; Scheffler et al., 2018). Parameters were:

<u>VASO:</u> voxel size = 0.82×0.82×1.5 *mm^3^*, TI1/TI2 = 1593/3780 ms, volume acquisition time (nulled and not-nulled combined) = 4872 ms, TE = 25 ms, GRAPPA acceleration factor = 3, phase Partial Fourier = 6/8, bandwidth 1064 Hz/px, and matrix size = 162 × 216 × 26. A variable flip angle scheme was used to minimize T1-related blurring (see Huber et al., 2017).

<u>bSSFP:</u> voxel size = 0.3×0.3×3 *mm^3^*, slice acquisition time = 3 s, TE = 4.79 ms, TR = 9.58 ms, no parallel imaging, no Partial Fourier, nominal flip angle = 35 deg, bandwidth 174 Hz/px, and matrix size 302×320×1. Prior to acquiring data for each slice, a series of RF pulses were applied to ensure that the MRI signal reached a steady state. To avoid the banding artifact of bSSFP images, we carefully adjusted the RF frequency to keep the region of interest in the middle of the passband off-resonance profile. A series of bSSFP images were collected with RF frequency shift (Δf) from -70 Hz to 70 Hz with a step of 10 Hz. The optimal Δf was determined from the off-resonance profile from the location of interest in M1 gray matter.

Both functional sequences had partial brain coverage. Imaging slabs were placed in the left hemisphere to cover the hand knob (Yousry et al., 1997) in the upper part of M1, i.e., Brodmann area (BA) 4a, in line with previous M1 laminar fMRI studies (Beckett et al., 2020; Chai et al., 2020; Guidi et al., 2016, 2020; Han et al., 2021; Huber et al., 2017; Persichetti et al., 2020; Shao et al., 2021). Slabs were placed perpendicularly to the cortical surface of the hand knob.

### 2.4 Experimental paradigm and procedures

VASO was chosen as the main sequence to address our research question due to its CBV-weighted contrast associated with superior spatial specificity, ideal for layer-dependent studies (Huber et al., 2017, 2019; Jin & Kim, 2008). The strong microvascular weighting of VASO results in inherently reduced CNR compared with gradient echo BOLD (Huber et al., 2019). In combination with a relatively weak proprioceptive stimulus (passive movement of a single finger with a small range of motion), this resulted in noisy activation maps. To provide additional evidence that deep layer signal is a robust feature of proprioceptive stimulation, we opted to scan 2 additional subjects (independent from VASO subjects) using bSSFP with extremely high in-plane resolution (0.3 mm). The spin echo-like contrast mechanism of bSSFP results in BOLD contrast with higher tissue specificity than conventional gradient echo BOLD (Báez-Yánez et al., 2017; Miller, 2012; Scheffler et al., 2018). It additionally benefits from minimal spatial distortions and blurring-effects because of its very short read-out time, which further improves the ability to resolve laminar activation (Liang et al., 2022; Miller, 2012). One subject was scanned twice across different days to assess the reliability of 2D-bSSFP activation maps (Figure S1). Prior to scanning, subjects practiced to reproduce the passive movement pattern during active movements of the right index finger and were instructed to completely relax during the passive condition. Movement frequency was guided visually by blinking text saying either ‘’you tap’’ for the active condition, ‘’machine tap’’ for the passive condition or ‘’relax’’, presented with an MRI compatible projector onto a semi-transparent screen (1024×768, 60Hz). The visual stimuli were programmed in MATLAB (Mathworks Inc.) using the Psychophysics Toolbox (Brainard, 1997). Subjects were asked to keep their fixations on the visual stimuli during the whole experiment, except in resting periods between runs. Each session consisted of an anatomical scan (T1-weighted MP2RAGE) followed by 2 functional runs per condition for VASO sessions and 3-4 runs per condition for bSSFP. Each run had a duration of 12 minutes for VASO and 6 minutes bSSFP. In active runs, 30 seconds of rest alternated with 30 seconds of active movement with the right index finger. Passive runs alternated between 30 seconds of rest and 30 seconds of passive movement where the device would move the right index finger during relaxation (Figure 1C). The order of conditions was counterbalanced across subjects to avoid order effects, and after finishing the runs of one condition, subjects were asked to slowly shift from the active to the passive device (or vice versa) while trying to minimize head motion. Head motion was restricted by custom-made bite bars.

A functional localizer run (12 minutes of active finger movements) using a gradient echo EPI BOLD sequence (voxel size: 0.75 × 0.75 × 0.86 *mm^3^*) was included at the end of VASO sessions for the purpose of ROI definition (see Methods Section 2.5.2).

### 2.5 Data analysis

#### 2.5.1 Preprocessing

The VASO data underwent standard preprocessing steps in line with previous laminar VASO studies (Finn et al., 2019; Huber et al., 2017). SS-SI-VASO produces two time-series in an alternating fashion, one where blood-signal is nulled (VASO time-series) and another without blood-nulling (BOLD time-series). Nulled and not-nulled images were motion corrected separately with the application of a spatial weighting mask to optimize the realignment around M1 using SPM12 (Functional Imaging Laboratory, University College London, UK). To correct for the influence of positive BOLD effect, which counteracts the negative CBV-response, dynamic division of nulled and not-nulled images was implemented after trial averaging as explained previously (Finn et al., 2019; Huber et al., 2014). Realigned nulled and not-nulled images were further used to compute an image with T1-weighted anatomical contrast (Huber et al., 2017), which was used as a “high quality reference image” for nonlinear registration of the MP2RAGE image to EPI-space. This co-registration step was performed by combining rigid, affine and non-linear (*SyN-algorithm*) transformations in ANTs (Avants et al., 2011).

The bSSFP data was preprocessed using AFNI (Cox, 1996), including removal of outliers (*3dToutcount* function), rigid body motion correction, brain masking, and coregistration of the T1 weighted anatomical volume to the mean functional image with localized pearson correlation as the cost function. Functional bSSFP images has minimal distortions, analogous to MP2RAGE, and high-quality coregistration was thus possible based on rigid and affine transformations only (Liang et al., 2022).

For both sequences, activation maps were computed voxel-wise as the average of condition-block timepoints minus the average of resting-block timepoints. This difference was converted to percent signal change by dividing by the average of resting-block timepoints. Note that the first two nulled and not-nulled volumes and the first four bSSFP volumes were discarded from each block to limit the influence of transition periods. Also, activation-induced CBV-increases result in negative VASO signal changes, and the corresponding activation maps were thus inverted.

#### 2.5.2 Region of interest (ROI) definition

We intended to assess the laminar pattern of passive movements within the same functional region as in previous laminar M1 studies, i.e., the part of the hand knob in area BA4a expressing a double peak response pattern during active movements. For VASO, ROIs were thus defined by first identifying the hand knob based on its omega-shaped anatomical landmark (Yousry et al., 1997). Then, VASO activation maps of the active condition were used to localize the area which most clearly expressed a double peak response pattern. Since VASO activation maps had relatively low CNR, independent localizer maps with BOLD contrast were used to confirm robust responses to index finger movements in the selected area. Connected ROIs spanning 1-2 slices and without holes were manually drawn in this area, with columnar widths roughly corresponding to that expected from individual finger representations in M1 (Huber et al., 2020). The same procedure was followed for bSSFP, except it was done on the single available slice.

For the subject that was scanned twice with 2D-bSSFP, we selected the session with strongest activation for the active condition to evaluate the response to passive movements. The other session was used to assess the robustness of activation maps across sessions (Figure S1).

Note that since data from the active condition was used to extract ROIs, we only statistically analyzed data from the passive condition. Laminar profiles corresponding to the active condition are still shown for reference, but are likely biased (Kriegeskorte et al., 2009) and should be interpreted as such.

#### 2.5.3 Laminar profiles and assignment of cortical layers to relative depths

For VASO, WM/GM and GM/CSF boundaries were drawn manually around ROIs based on MP2RAGE images and used to compute equidistant depth maps using LAYNII (Huber et al., 2021). This was done on an upsampled grid with 0.2 mm in-plane resolution. The same procedure was followed bSSFP, except that upsampling was not required due to the inherently high in-plane resolution of this sequence. Laminar profiles with 18 bins were generated for each condition by computing the average percent signal change across ROI-voxels for each depth bin. Profiles were extracted from all ROI voxels with no thresholding being applied.

Layer Va’s approximate depth and width were inferred from laminar thickness estimates outlined in Palomero-Gallagher & Zilles, (2019) and Zilles & Amunts, (2012). This depth co-localized with the dip between deep and superficial peaks in the active condition’s group-level VASO profile, and the T1-profile plateau, previously used as M1 layer Va landmarks (Huber et al., 2017) (see Figure S2 for details).

#### 2.5.4 Statistical analysis

Cluster-based permutation testing (Nichols & Holmes, 2003) was employed for determining the significance of bins in passive laminar profiles, which tackles the issue of multiple comparisons. Testing was performed across subjects for VASO (group-level) and across trials for bSSFP (subject-level). For each permutation, passive laminar profiles were randomly inverted with a minus sign since under the null-hypothesis, all profiles could equally likely have come from the contrast “rest>passive” as from the contrast “passive>rest”. A two-sided p-value was then calculated for each bin across resulting profiles using one-sample t-test. The maximum t-value sum observed for a cluster (range of consecutive bins with uncorrected p-values < *0*.*05*) was recorded for each permutation. This was repeated in 100.000 permutations, yielding a null distribution from which the p-value could be derived for the t-value sum of a given cluster in the measured data. This method tests the significance of a cluster as a whole, but significance of individual bins within the cluster cannot be inferred (Nichols & Holmes, 2003). Thus, to evaluate significance of deep layer bins in isolation (without contribution from bins in superficial layers), p-values were also derived for sub-clusters that only contained deep layer bins (between the GM/WM boundary and layer Vb/Va boundary).

The significance level for all statistical tests was set at *α*=0.05.

### 2.6 Data and code availability

The raw data will be available for download via figshare (https://doi.org/10.6084/m9.figshare.23834838). Analysis code will be made publicly available on GitHub (https://github.com/LasseKnudsen1/7TactiveVSpassiveStudy/tree/main).

## 3. Results

To test whether human M1 deep layer responses are evoked by proprioception, laminar profiles were extracted from ROIs containing the characteristic double peak response pattern for active movements (see Methods Section 2.5.2). Figure 2A depicts the raw VASO activation maps of an example subject for each condition. The active map has robust activation in the expected part of the hand knob (BA4a), but maps are obviously noisy (with many false positives) and the double peak feature is not clearly visible (Figure 2A, upper middle). Therefore, we generated within-layer smoothed activation maps (Huber et al., 2021), from which the double peak pattern more clearly emerged in the active tapping condition (Figure 2A, upper right). The smoothed map for the passive condition also shows superficial and deep activations in similar locations (as indicated by arrows in Figure 2A, lower right) but were weaker compared with the active tapping condition. Figure 2B shows the group-level VASO profile associated with active finger movements. Importantly, the shape and response magnitude of this profile are biased toward the ROI-selection criteria and should be interpreted accordingly (Kriegeskorte et al., 2009). It also shows the group-level passive laminar profile derived from independent ROI-selection criteria (based on activations in the active condition) and thus unbiased (for individual subject’s profiles, see Figure S3). Significant clusters of activation were found in both the superficial (p = 0.047) and deep layers (p = 0.016) (shown as horizontal black lines in Figure 2B; numbers denote a significant sub-cluster of deep layer bins, p = 0.027). Interestingly, the deep layer peak appears to be slightly displaced towards layer Va during passive movements compared to active movements (Figure 2B).

**Figure 2.**
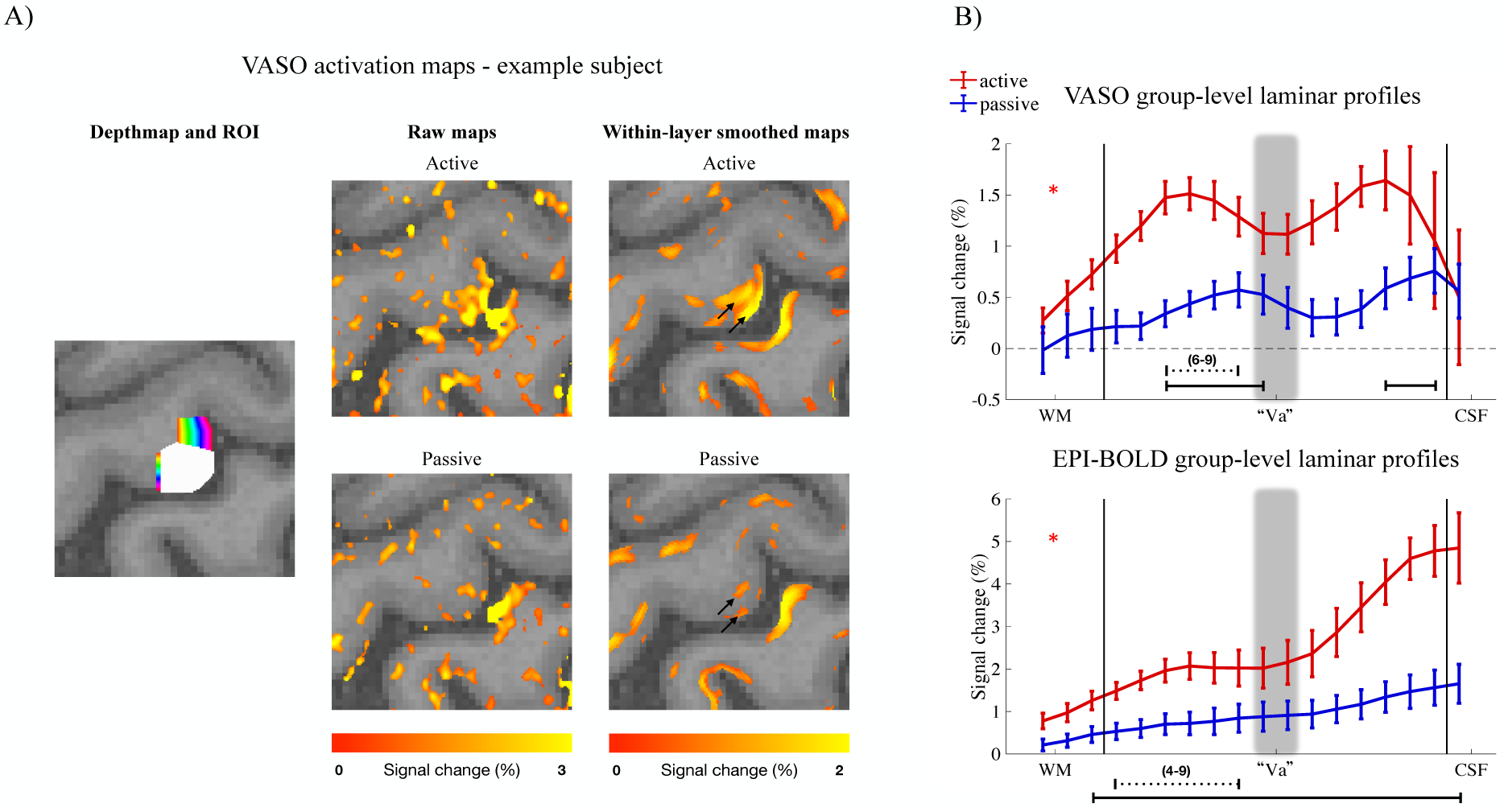
Evaluation of deep layer signal during passive movements using VASO **A)** The first column shows the depth map and ROI used to extract VASO and EPI-BOLD laminar profiles for this example subject. The second columns show raw percent signal change maps for each condition and the third column shows corresponding maps after within-layer smoothing. Arrows point to indications of the double peak feature. Maps are thresholded at t=1.5. **B)** Laminar group-level profiles for each condition, both for VASO and BOLD. Asterisks (red) denote circular profiles for the active (red) condition. Solid horizontal lines illustrate significant clusters as evaluated by permutation tests. Dotted lines and corresponding numbers denote bins (counting from left to right) contained in a significant deep layer sub-cluster (Methods section 2.5.4). Solid vertical lines illustrate WM/GM and GM/CSF boundaries. The blurry gray shaded area denotes the approximate position of layer Va. Error bars refer to standard error of the mean across subjects.

Activation maps (Figure S4A) and laminar profiles (Figure 2B, Figure S3) were similarly generated from the not-nulled timeseries of the VASO-sequence (referred to here as EPI-BOLD to avoid confusion with the BOLD-contrast of bSSFP). In line with VASO results, significant activation was found in both superficial and deep layers (all GM bins were above threshold resulting in a single large cluster shown with the horizontal black line in Figure 2B, p = 0.002; numbers denote significant sub-cluster of deep-layer bins, p = 0.023). However, maps and profiles were characterized by a bias towards superficial layers, and no clear double peak was observed at the group-level in either condition. This is expected due to the sensitivity of gradient echo EPI-BOLD towards draining veins, which displace the signal from local neuronal activity and thus limit its laminar specificity (Menon, 2012; Turner, 2002). In an attempt to account for the signal dispersion originating from intracortical veins, spatial deconvolution (Markuerkiaga et al., 2016; Marquardt et al., 2018) was applied to the profiles in a supplementary analysis (Figure S4B). After deconvolution, the profiles began to resemble the corresponding VASO profiles (low superficial bias and indications of double peaks), known to be less influenced by macrovascular contamination. Nevertheless, considering the limitations of the applied correction strategy (see Figure S4B), the corrected profiles should be interpreted with caution.

The activation maps for bSSFP are shown in Figure 3A. Only one of the subjects had a clear double peak response within the same area in both conditions, but both subjects appear to have robust deep layer activation in the passive tapping condition where activation of the active condition mostly resembled the double peak feature (denoted by arrows). This is supported by both subjects’ laminar profiles (Figure 3B) showing significant sub-clusters of activation in the deep layers (p = 0.005 and p = 0.001 for Subject 1 and Subject 2, respectively). Due to bSSFP being relatively insensitive to T2* contributions, it is expected to have less bias towards large veins than gradient echo EPI (Báez-Yánez et al., 2017; Liang et al., 2022; Miller, 2012; Scheffler et al., 2018). Nevertheless, as visible in activation maps, we did observe strong signals towards CSF, which likely reflects high sensitivity of bSSFP-BOLD to intravascular signals and inflow effects (Liang et al., 2022). To minimize the effect of large vessels, voxels with percent signal change exceeding 4% in the across-condition averaged bSSFP map were excluded from the map of each condition before generating laminar profiles (uncorrected profiles are shown in Figure S5).

**Figure 3.**
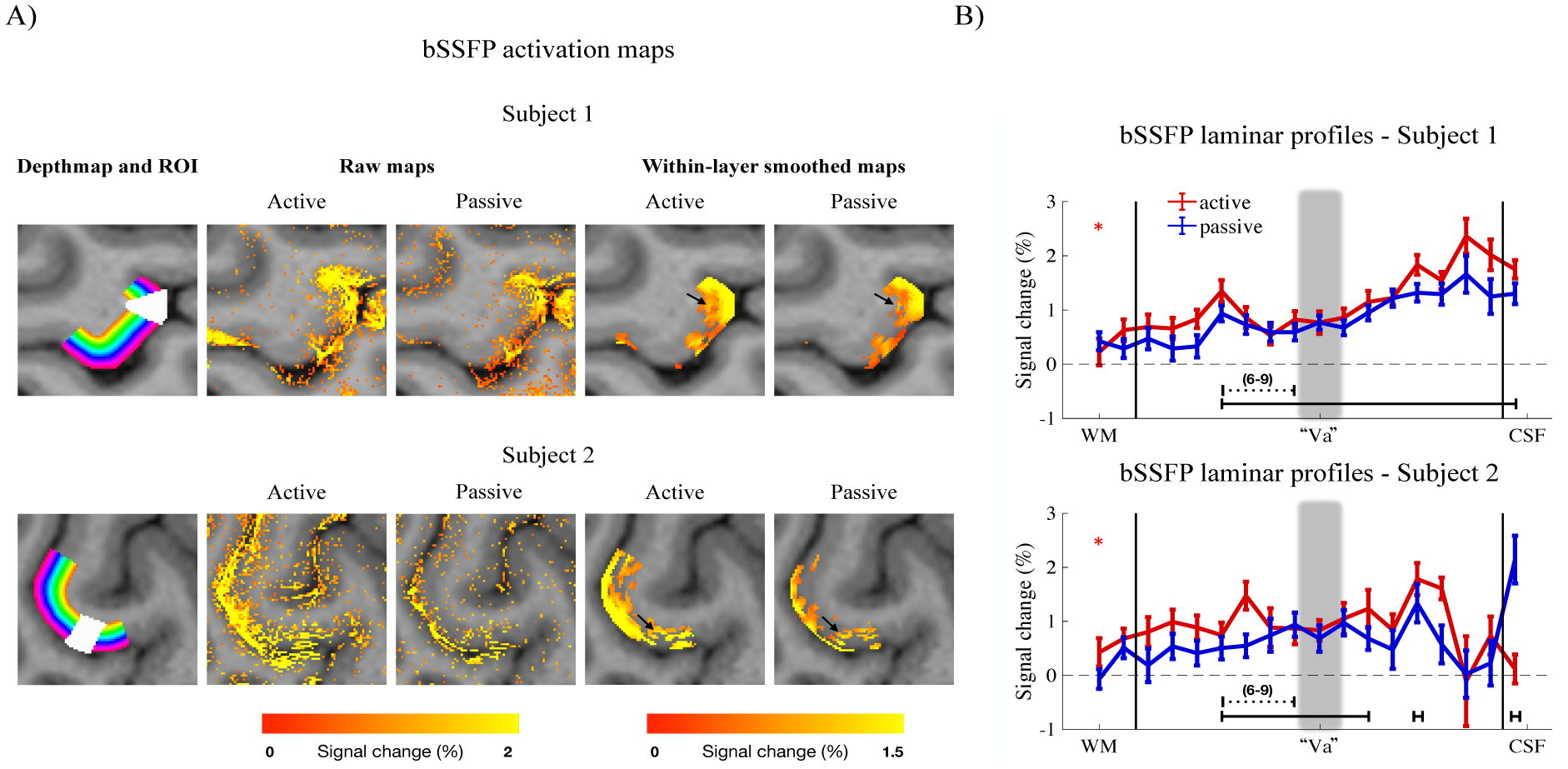
Evaluation of deep layer signal during passive movements using bSSFP **A)** The first column shows the depth map and ROI used to extract bSSFP laminar profiles for each subject. The next columns show raw percent signal change maps for each condition and the last columns show corresponding maps after within-layer smoothing. Arrows point to locations of robust deep layer activation. Raw maps are thresholded at t=2 and smoothed maps at t=3. **B)** Subject-level laminar profiles for each condition. Asterisks (red) denote circular profiles for the active (red) condition. Solid horizontal lines illustrate significant clusters as evaluated by permutation tests. Dotted lines and corresponding numbers denote bins (counting from left to right) contained in a significant deep layer sub-cluster (Methods section 2.5.4). Solid vertical lines illustrate WM/GM and GM/CSF boundaries. The blurry gray shaded area denotes the approximate position of layer Va. Error bars refer to standard error of the mean across trials.

## 4. Discussion

In the present study, we used submillimeter resolution VASO and bSSFP at 7T to demonstrate significant deep layer activation in human M1 in response to proprioceptive stimulation. In the following sections, we first relate this finding to the existing literature, followed by interpretations of the M1 deep layer signal in the context of previous laminar fMRI studies and the active inference account of motor control. The last section discusses study limitations.

### 4.1 Passive movements activate both superficial and deep layers of M1

Multiple studies have investigated the M1 laminar activation pattern associated with active finger movements. A consistent finding is the characteristic double peak response within the hand knob of BA4a (Beckett et al., 2020; Chai et al., 2020; Guidi et al., 2016, 2020; Huber et al., 2017; Persichetti et al., 2020; Shao et al., 2021). This could similarly be observed in the present study, both in VASO (Figure 2 and Figure S3) and bSSFP (Figure 3) experiments. The novel finding of the present study is that deep layers also activate in response to passive movement, i.e., a task in which voluntary motor aspects are presumably absent. Specifically, from the group-level laminar profiles, our VASO results revealed significant activation in both superficial and deep layers during proprioceptive stimulation (passive condition in Figure 2), which was supported by EPI-BOLD results. In addition, both bSSFP subjects displayed robust deep layer responses visible in passive activation maps (Figure 3A) and significantly activated bins in deep layers of the corresponding laminar profiles (Figure 3B). Accordingly, our findings suggest a direct proprioceptive influence on M1 deep layer signals.

To our knowledge, M1 activation in response to proprioceptive stimulation has not been studied at the human laminar level previously, but, consistent with our findings, numerous studies have established strong M1 involvement in relation to proprioceptive perception. For example, proprioceptive sensations can be induced by tendon vibration which elicits an illusion that the corresponding limb is moving. M1 has been shown to be the most strongly activated area during such illusions, despite the fact that subjects did not actually move the limb nor intended to do so. Activation was only found when the tendon was vibrated at frequencies that did not elicit the illusion, thereby excluding tactile influence (Naito, 2004). Similarly, subjects experience kinesthetic illusions when attempting to move a limb during experimentally induced paralysis, even when afferent somatosensory signals are blocked (Gandevia et al., 2006; Proske & Gandevia, 2012; Smith et al., 2009; Walsh et al., 2013). Another line of evidence comes from the observation that proprioceptive stimulation significantly increases M1 activation (Blatow et al., 2011; Ciccarelli et al., 2005; Guzzetta et al., 2007; Jaeger et al., 2014; Mehta et al., 2012; Weiller et al., 1996), and modulates M1 excitability (Carel et al., 2000; Lewis & Byblow, 2004; Onishi, 2018). These findings suggest an important role of M1 in the proprioceptive hierarchy, and demonstrate that voluntary movement components are not a prerequisite for M1 activation. The present study contributes to the literature on humans by showing that both superficial and deep layers are involved. This is consistent with non-human primate electrophysiological studies showing that a large proportion of M1 cells are responsive to proprioceptive stimulation (see Naito 2004, and references therein), and that such cells are found in both superficial and deep layers, including corticospinal pyramidal tract neurons (Lemon & Porter, 1976; Wiesendanger, 1973).

### 4.2 Interpretation of the deep layer signal in M1

Huber et al. (2017) suggested that the peak in superficial layers mainly reflects corticocortical input from somatosensory (tactile and proprioception) and premotor areas. This interpretation was informed by anatomical tracer studies in rodents (Mao et al., 2011; Weiler et al., 2008, see also Figure 1A) and was supported by their resting state connectivity results. Furthermore, tactile stimulation was shown to affect superficial layers only, and in a subsequent study, Persichetti et al., (2020) showed similar findings for imagined movements believed to entail substantial input from premotor areas. In other words, tasks devoid of corticospinal motor output appear to lack deep layer signal changes, whereas active movements, assumed to involve motor output, are linked to deep layer activation. When factoring in that corticospinal neurons are selectively located in layer Vb (Figure 1A), it is reasonable that the double peak has commonly been interpreted as premotor/somatosensory input in superficial layers, and voluntary-movement-related corticospinal output in deep layers (Beckett et al., 2020; Chai et al., 2020; Dumoulin, 2017; Huber et al., 2017; Larkum et al., 2018; McColgan et al., 2020; Persichetti et al., 2020; Shao et al., 2021). Our observation of significant deep layer activation during passive movements is indicative of a concomitant proprioceptive contribution to the deep layer signal. Considering that M1 strongly relies on proprioceptive state information for proper generation of motor commands (Hatsopoulos & Suminski, 2011; Omrani et al., 2017; Proske & Gandevia, 2012), the observed deep layer signal may reflect delivery of such proprioceptive information to pyramidal tract cells in layer Vb to inform potential motor output. Based on the model proposed in Figure 1A, such signals mainly arrive in superficial layers where they undergo preprocessing, before being relayed to deep neurons through the strong superficial-to-deep intra-columnar connection (Bastos et al., 2012; Weiler et al., 2008). On the other hand, M1 has been shown to be more than just a passive recipient of proprioceptive information; it actively contributes to the generation of proprioceptive perceptions (Gandevia et al., 2006; Naito, 2004; Nudo et al., 2000). Thus, deep output layers may be actively involved in shaping proprioceptive sensations, as opposed to merely using it to guide output.

In view of the increasing prominence of hierarchical Bayesian theories of brain function and their inherent link to laminar circuitry, we propose an alternative interpretation of the observed deep layer signal. This is based on the active inference framework (Friston, 2010), which is particularly relevant to the sensorimotor system. According to the canonical microcircuit underlying active inference (Bastos et al., 2012), proprioceptive stimulation should strongly activate superficial layers of M1 since neuronal units at this depth are the targets of driving feedforward proprioceptive prediction errors which continuously update expectations about the current proprioceptive state. The potent superficial-to-deep intracolumnar pathway conveys these expectations onto deep layer prediction units, which is accompanied by feedback connections from premotor cortex. Thus, deep layer units integrate extensive synaptic input, which in itself may give rise to deep layer fMRI signals (Logothetis, 2008). Furthermore, on the basis of this input, proprioceptive predictions are generated and transmitted to S1 and thalamic somatosensory nuclei to inform proprioceptive perceptions (*perceptual inference* component of active inference). Within this framework, corticospinal output constitutes these same proprioceptive predictions directed to lower motor neurons in the spinal cord (the distinction between motor output and proprioception, as in Figure 1B, thus becomes meaningless as motor output and proprioception are two sides of the same coin). Cortical predictions about the current proprioceptive state are compared with incoming data from the muscular Ia stretch receptors. Any disparity between prediction and data yields a proprioceptive prediction error which activates the motor neuron until the prediction is realized in the subsequent movement (*action* component of active inference, for details, see Adams et al., 2013; Barrett & Simmons, 2015; Bastos et al., 2012; Friston, 2011; Shipp et al., 2013). Consequently, another possible explanation for the observed deep layer signal is corticospinal drive being present even during passive movements; if the afferent data from the externally induced movement is not matched by a corticospinal proprioceptive prediction, a prediction error ensues leading to a counter movement (akin to the classical stretch reflex, Lidell & Sherrington, 1924). Thus, in order for subjects to be able to relax, M1 may continuously transmit corticospinal drive in the form of predictions about proprioceptive consequences of the passive movement. This interpretation is consistent with the observation that pyramidal tract neurons in layer Vb fire in response to passive movements (Lemon & Porter, 1976; Wiesendanger, 1973). Nevertheless, we cannot exclude that the observed deep layer signal originated from non-pyramidal tract cells and more evidence for corticospinal drive during passive movements is necessary to support this explanation.

Interestingly, the deep peak for the passive condition appears slightly shifted towards layer Va compared to the active condition (Figure 2B). Layer Va is the main connection between M1 and S1 (Hooks, 2017; Mao et al., 2011), whereas layer Vb contains all corticospinal cells. Thus, one possible explanation is that the proprioceptive predictions transmitted to S1, mediating *perceptual inference*, dominates during passive movements whereas corticospinal drive exerts more influence during *action* (Adams et al., 2013). The shifted peak may also be related to the hypothesis by Hooks et al., (2013) that two different subpopulations of pyramidal tracts neurons exist in M1; one in the superficial end of layer Vb, mainly influenced by somatosensory aspects (referred to as the sensorimotor output channel), and another in the deeper end of layer Vb, influenced more by cognitive facets such as intention to move (referred to as frontomotor output channel). These interpretations are not mutually exclusive. However, both of them are rather speculative, and further studies are needed to corroborate whether the shift is a robust feature distinguishing laminar profiles of active and passive finger movements.

### 4.3 Limitations

Our conclusion that proprioceptive stimulation activates deep layers of M1 is partly based on the significant cluster of activation in the deep end of the group-level laminar VASO profile shown in Figure 2B. However, large variability was observed in the laminar activation pattern across subjects (Figure S3). For some subjects, the deep layer percent signal change was even zero or below. The same conclusion is drawn based on the BOLD-EPI and bSSFP data with higher sensitivity, which compensates for this limitation to some extent. Nevertheless, we do acknowledge that our findings would be more convincing if robust laminar patterns of VASO activation could be demonstrated across subjects for passive movements, as Huber et al. (2017) did for active movements. Part of the variability across subjects is likely explained by the previously mentioned CNR limitation of VASO. Also, in our finger movement task, only one finger was moved, and the range of motion was small due to the passive movement device’s constraints. To increase response magnitudes (and thus CNR), it would be interesting to replicate the experiment of the present study using a passive movement device that tolerates multi-finger movements and generates a wider range of motion.

It should finally be noted that our findings are only valid if subjects relaxed their fingers during passive movements, which could have been confirmed through EMG recordings. Although we did not obtain such recordings, previous research has demonstrated that subjects generally have no difficulty relaxing their muscles during passive movements (Ciccarelli et al., 2005; Mima et al., 1999; Thickbroom et al., 2003).

## 5. Conclusion

We used 7T laminar fMRI to investigate the laminar pattern of activation in M1 in response to passive finger movements. Both VASO and bSSFP displayed significant activation in both superficial and deep layers. This finding was interpreted in the light of previous laminar fMRI research and the framework of active inference. We conclude that M1 deep layers constitute an important component in proprioceptive circuits and that a non-negligible proportion of the M1 deep layer activation likely is due to proprioceptive influence. We believe this study enhances our understanding of the laminar circuitry in human M1, thereby facilitating interpretations of future laminar research and contributing to the development of theoretical frameworks of motor control and somatosensory perception.

## Acknowledgements

The authors would like to thank all subjects for their participation. We would further like to thank Wu Chen and Yanhui Fu (Institute of Biophysics, Beijing, China) for technical assistance, as well as Chengwen Liu and Chencan Qian (Institute of Biophysics, Beijing, China) for guidance on analysis. We thank Christopher J. Bailey (Aarhus University Hospital) for guidance throughout the project and development of the passive movement device. This work was supported by Beijing Natural Science Foundation (Z210009), STI2030-Major Projects (2022ZD0204800, 2022ZD0211900, and 2021ZD0204200), National Key R&D Program of China (2022YFB4700101), the National Natural Science Foundation of China (32070987, 31722025, and 31930053), the Strategic Priority Research Program of Chinese Academy of Sciences (XDB37030303), and a grant from Sino-Danish Center (SDC).

## Credit authorship contribution statement

**Lasse Knudsen:** Conceptualization, Methodology, Formal analysis, Investigation, Writing - Original Draft, Writing - Review & Editing, Visualization, Funding acquisition

**Fanhua Guo:** Methodology, Formal analysis, Investigation, Writing - Review & Editing, Visualization

**Jiepin Huang:** Methodology, Investigation, Writing - Review & Editing

**Jakob U. Blicher:** Conceptualization, Methodology, Writing - Review & Editing, Supervision, Project administration, Funding acquisition

**Torben E. Lund:** Conceptualization, Methodology, Writing - Review & Editing

**Yan Zhou:** Conceptualization, Methodology, Writing - Review & Editing, Project administration, Funding acquisition

**Peng Zhang:** Conceptualization, Methodology, Writing - Review & Editing, Supervision, Project administration, Funding acquisition

**Yan Yang:** Conceptualization, Methodology, Writing - Review & Editing, Supervision, Project administration, Funding acquisition

