## Supplementary figures for "The laminar pattern of proprioceptive activation in human primary motor cortex"

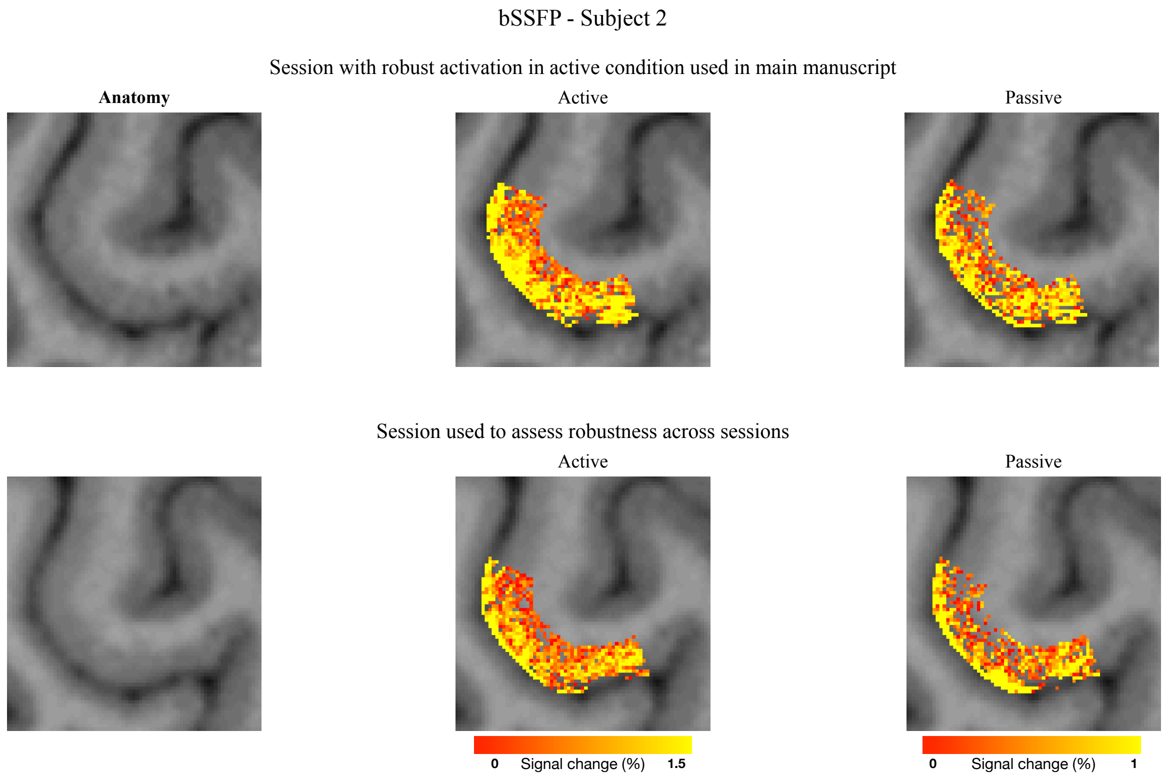


**Figure S1.** Across-session robustness of bSSFP activation maps. This subject was scanned with bSSFP in two different experimental sessions separated by several days. The first column shows anatomy for each session. The second and third columns show masked activation maps for active and passive conditions, respectively. Visual inspection suggests similar patterns of activation, i.e., roughly the same areas light up in both sessions, but the magnitude of activation was relatively weak for both conditions in the second session (lower panel).

Laminar profiles of T1-values - data from VASO experiment

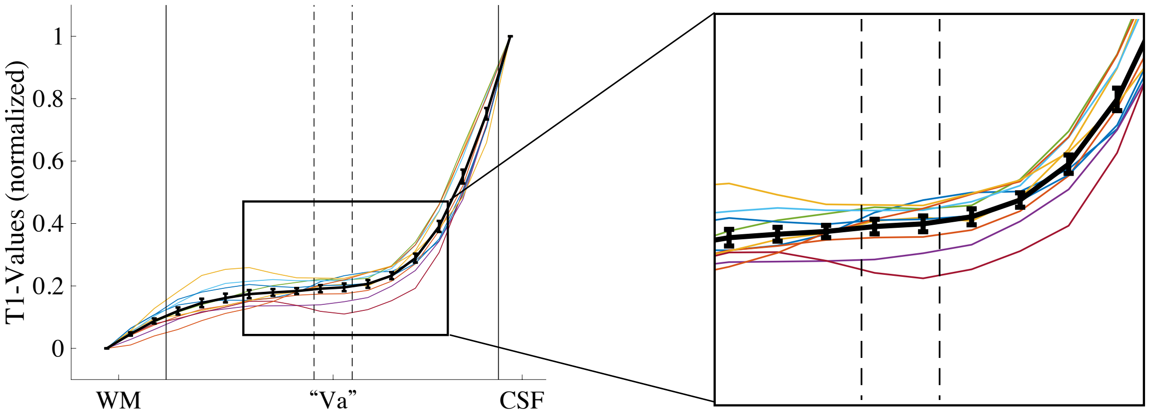


**Figure S2.** Validation of layer Va positioning using T1 profiles from the VASO experiment. This figure shows the group-mean laminar profile (black) and subject-level profiles (colored) of T1-values (derived from MP2RAGE, Marques et al., 2010) extracted from ROIs in the VASO experiment. Dashed vertical lines represent the expected Va position solely based on relative thickness estimates of each layer outlined in (Palomero-Gallagher & Zilles, 2019; Zilles & Amunts, 2012). Huber et al., (2017) suggested that the approximate depth of layer Va can be assigned to the location where the group-level VASO laminar profile of the active condition has a dip between superficial and deep peaks. In this study, this landmark co-localized with the estimated position based on relative thickness estimates. Huber et al. (2017) further showed that the plateau of T1-laminar profiles represents another landmark for Va. There is no clear plateau in the group-level T1 profile shown here, but it seems to flatten in the location around the dashed lines. Furthermore, as visible in the zoomed view, some subjects had identifiable plateaus, and these appear to fall approximately within the dashed lines.

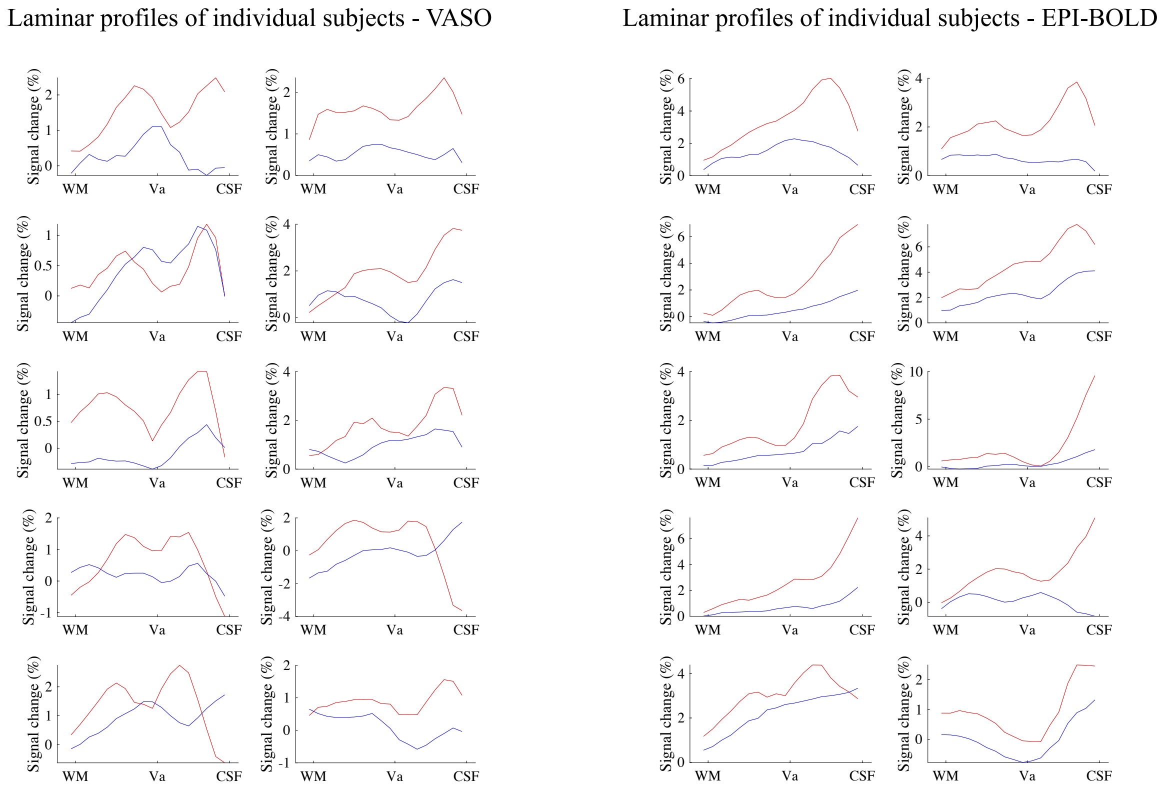


**Figure S3.** VASO and EPI-BOLD laminar profiles of individual subjects. Note that all active profiles (red) are non-independent with respect to ROI definition, and thus potentially biased. Passive profiles (blue) are independent and unbiased.

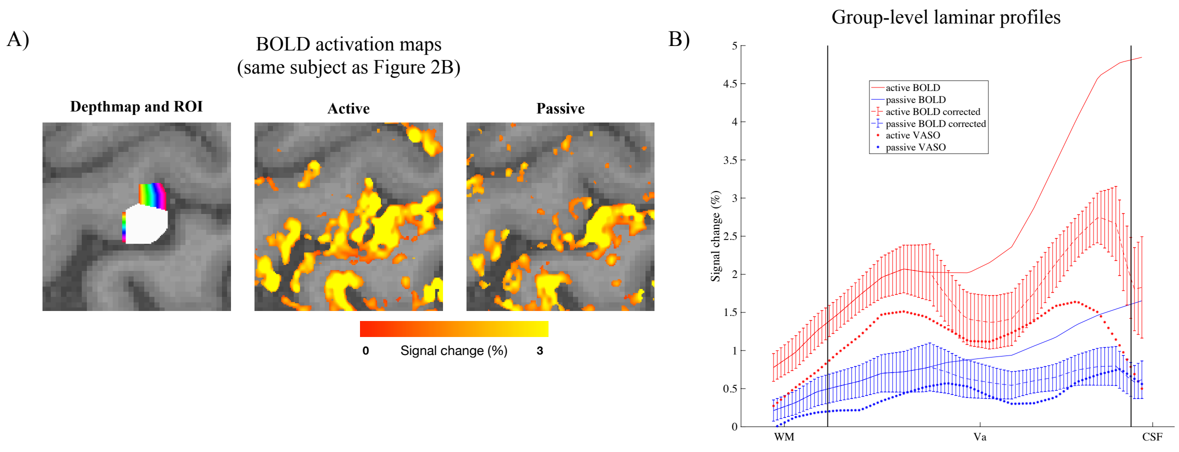


**Figure S4. A)** EPI-BOLD activation maps corresponding to those shown for VASO in Figure 2A (same subject). Maps are thresholded at t=2.5 **B)** The EPI-BOLD laminar profiles from Figure 2B (solid profiles in this plot) are characterized by a positive gradient from deep to superficial layers, and no clear double peak is observed. This is expected when using gradient-echo BOLD with EPI-readout due to its strong sensitivity towards pial veins and intracortical veins that drain blood towards the cortical surface (Turner, 2002). Notice the absence of the gradient in VASO profiles (asterisk profiles), known to be relatively insensitive to veins. It should be mentioned that our study mainly cares about deep layer responses which are expected to be relatively unaffected by interlaminar drainage due to its unidirectional nature. Still, we tried to correct for the effect of signal drainage on laminar profiles using spatial deconvolution (Markuerkiaga et al., 2016; Marquardt et al., 2018). Briefly, profiles were first interpolated to 100 equally spaced depth bins, then each bin was assigned to a layer based on relative layer thicknesses (derived from Palomero-Gallagher and Zilles (2019); Zilles & Amunts (2012)), and the corrected signal in each bin was then given as the original signal minus the contribution from the layers below (see analysis code: https://github.com/LasseKnudsen1/7TactiveVSpassiveStudy/tree/main). After this correction scheme, the superficial bias was largely gone and double peaks emerged in the profiles of both conditions. However, the corrected profiles should not be overinterpreted. The deconvolution relies on accurate weighting parameters describing the contribution of signal in each bin that originate from the bins below. As in Marquardt et al., (2018), we used the parameters derived from the visual cortex (Markuerkiaga et al., 2016), which may not be readily applicable in M1 due to vascular differences, and due to a different laminar organization (layer IV is for example largely absent in M1). Also, this approach assumes identical drainage between all bins within a layer which is likely inaccurate. It does seem promising, though, that the corrected profiles are reminiscent of the corresponding VASO profiles, which was also observed in Marquardt et al., (2018).

A)

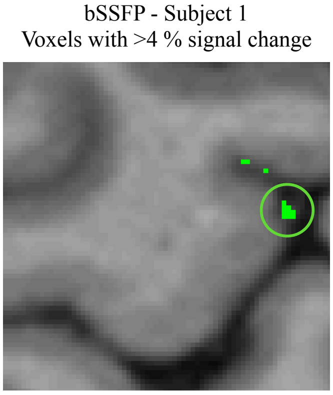


B)

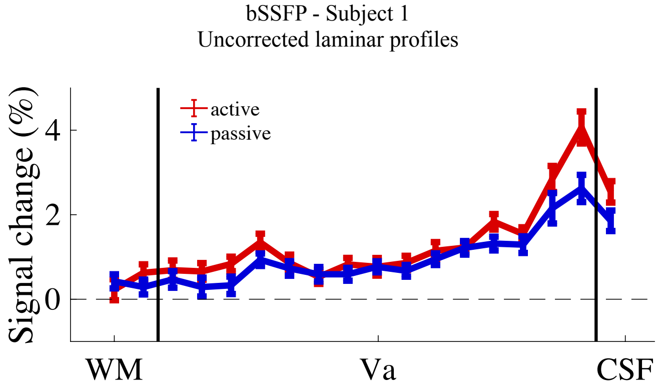


**Figure S5. A)** To reduce the influence of large veins on bSSFP laminar profiles, we excluded voxels where the percent signal change of the across-condition averaged activation map exceeded 4 % (absolute value) as suggested by (Liang et al., 2022; Liu et al., 2020). Green voxels have absolute percent signal changes larger than 4 %. Those highlighted by the ring were contained within the ROI. These voxels are located towards the cortical surface as expected if the strong response is dominated by macrovasculature. **B)** Same laminar profiles as in Figure 3B, but without exclusion of the voxels highlighted in A). The overall shape of the profiles is largely unchanged, but the macrovascular-induced bias towards the surface is more pronounced. Data is not shown for the second subject as no voxels in the ROI exceeded 4 %.
